# A single-cell eQTL atlas of the human cerebellum reveals vulnerability of oligodendrocytes in essential tremor

**DOI:** 10.1101/2024.05.22.595233

**Authors:** Charles-Etienne Castonguay, Farah Aboasali, Miranda Medeiros, Théodore Becret, Zoe Schmilovich, Anouar Khayachi, Alex Rajput, Patrick A. Dion, Guy A Rouleau

## Abstract

Essential tremor (ET) is a movement disorder characterized by an upper-limb postural and action tremor. It is one of the most common neurological disorders, affecting 1% of the worldwide population. Despite strong evidence for genetic factors driving the aetiology of ET, the underlying pathophysiology remains poorly understood. To understand the effects of genetic risk factors in ET on the cerebellum, the brain region thought to be affected by the disease, we built a population-scale single-cell atlas of the human cerebellar cortex comprised of over 1 million cells from 109 individuals. Using single-cell expression quantitative trait loci and mendelian randomization, we found evidence of ET-associated variants in the *BACE2* locus causally linked to its downregulation in cerebellar oligodendrocytes. We highlight a genetically vulnerable population of *BACE2-*expressing immature oligodendrocytes, suggestive of demyelination. We also find dysfunctional processes affecting interactions between Golgi cells, Purkinje layer interneurons, and oligodendrocytes in ET. Our study suggests a crucial role for cerebellar oligodendrocytes in the pathogenesis of ET.

## Main

Genome-wide association studies (GWAS) have identified thousands of common variants associated with neurological and psychiatric disorders. The majority of these variants are found in non-coding regions of the genome. Understanding the effects of these variants remains difficult. It is hypothesized that these variants affect the transcription of nearby genes^1^. This led to the identification of expression quantitative trait loci (eQTLs), based on the use of population-scale bulk-RNA sequencing data. Recently, with the advent of single-cell sequencing technologies, discovery of cell-type specific eQTLs (scQTLs) have been a major focus of study^2–5^. In the brain, scQTLs have been found to colocalize with disease-associated variants in a cell-specific manner^2^. Although useful, recent brain scQTLs studies have, for the most part, used single-cell data aggregated across multiple cortical regions^2^. This neglects the heterogeneity of neurons and glia across different brain regions^6,7^. Seeking to investigate area specific scQTLs, we focused on the cerebellar cortex, a brain region implicated in many neurological and psychiatric diseases^8,9^ ^10–13^.

The cerebellum is a major structure of the hindbrain, crucial for the integration of proprioceptive, sensory, and cognitive information. It is composed of a tightly regulated neuronal network centered around Purkinje cells. These neurons receive proprioceptive information via thousands of granule cell projections onto their elaborate dendritic trees in the molecular layer. Purkinje cell activity is also regulated by other interneurons found in the molecular and Purkinje layer of the cortex. Purkinje cells then form the sole output signal of the cerebellum, projecting onto the dentate nucleus, thalamus, and cortical motor areas. Proprioceptive information is then integrated to coordinate motor control during movements.

Clinical and functional studies have highlighted the role of the cerebellum in neurodegenerative disorders such as amyotrophic lateral sclerosis (ALS)^14^, Parkinson’s disease (PD)^9,10^ and Alzheimer’s disease (AD)^11^. Although cerebellar atrophy has been observed in these disorders^15,16^, it is thought that cerebellar involvement might only occur later on in disease progression, especially in AD^17,18^. This is in line with infrequent cerebellar tau accumulations^19^ or Beta-amyloid deposits in AD^18^. The cerebellum is therefore postulated to be protected early on in neurodegenerative disorders. Unexpectedly, some cognitive functions attributed to the cerebellum (‘dysmetria of thought’) also seem to be affected in these diseases and others such as schizophrenia (SCZ)^13^.

Essential tremor (ET) causes uncontrollable postural and action tremors in about 1-5% of the world’s population and is one of the most common neurological disorders^20–22^. Although genetic and functional studies have highlighted the importance of the cerebellum in the disease, causal cell-types for ET remain mostly unknown^23–25^. It is thought that degeneration and loss of Purkinje cells is a hallmark of the disease, but this finding remains controversial^26–29^. Moreover, other populations, such as cerebellar interneurons have yet to be characterized in ET. Studying how genetic variation is regulated in the cerebellum could lead us reveal the factors underlying cerebellar vulnerability in ET but also its relative resilience in other neurodegenerative diseases.

In this study, we used population-scale single-cell sequencing coupled with genotyping to identify scQTLs in the cerebellar cortex. We find that scQTLs in the cerebellum share similar properties with previously identified scQTLs in the cerebral cortex. Using a mendelian randomization approach, we used our scQTL atlas to uncover causal associations between SNPs and gene expression for multiple neurological and psychiatric traits in the cerebellum. Focusing on ET, thought to be a cerebellar disease, we found that top genome-wide significant variants in ET were causal scQTLs for *BACE2* downregulation in oligodendrocytes. We found that a specific subpopulation of *BACE2*-expressing immature oligodendrocytes was driving disease-gene enrichment in ET. This subset was disproportionately more affected than other oligodendrocytes when comparing ET and healthy cells, displaying changes of mRNAs related to axonal and synaptic homoeostasis. Finally, using independent cohorts of single-cell and bulk-RNA sequencing of ET patients, we replicate findings concerning the selective vulnerability of oligodendrocytes in the disease. Using our atlas of the human cerebellum, we establish the first causal relationship between oligodendrocytes and ET.

## Results

### A single-cell atlas of the human cerebellum

We used split-pool ligation-based transcriptome sequencing (SPLiT-seq)^30^ to sequence a total of 1.08 million nuclei from the cerebellar cortex of 109 individuals (Figure 1A). Our cohort was comprised of 16 ET patients as well as 93 healthy controls (Figure 1A-C). We were able to retrieve, on average, 9,250 nuclei per sample, 3,486 unique molecular identifiers (UMIs) and 1,361 genes per nuclei.

**Figure 1.**
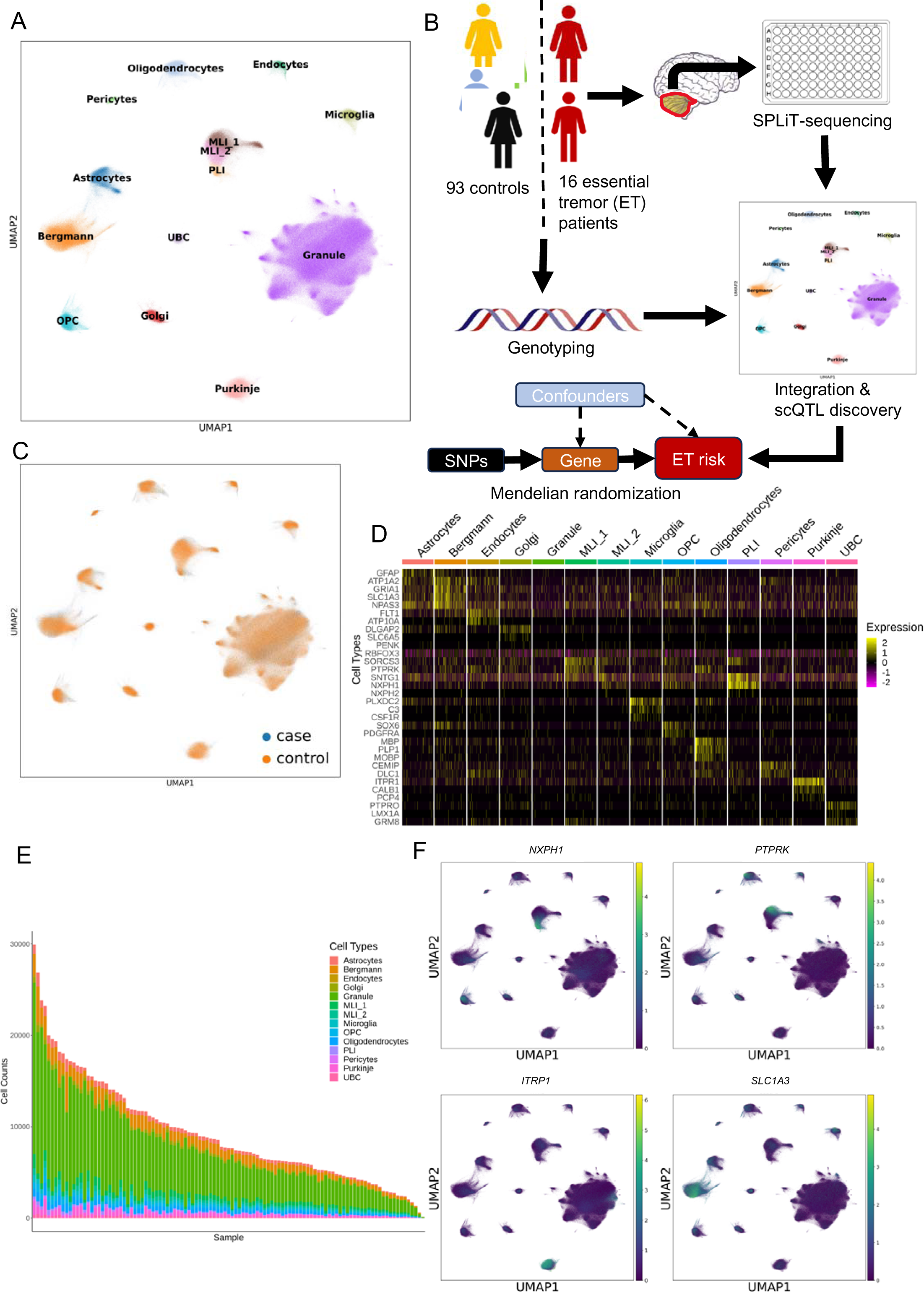
Characterization of a population-scale dataset of cerebellar nuclei from 109 individuals. UMAP clustering of 1.08M nuclei from 109 patients across 14 cell clusters. **B.** Experimental design for scQT discovery and identification of causal SNP-gene associations in the human cerebellum using mendelian randomization. **C.** UMAP clustering and integration of cerebellar cells across case-control status. **D.** Heatma of cerebellar cell-type markers across all 14 clusters. Colour denotes normalized expression counts. **E.** B plot of cell counts across 109 donors. Colour denotes cell-types. Note the overrepresentation of Granule cel in all samples. **F.** UMAP clustering with cell marker expression for PLI (*NPXH1*), MLI_1 (*PTPRK),* Purkin cells (*ITPR1)* and Bergmann glia (*SLC1A3)*. Scale bar represents normalized counts. Legend: MLI_ Molecular layer interneuron 1; MLI_2: Molecular layer interneuron 2; OPC: Oligodendrocyte progenitor cell PLI: Purkinje layer interneuron; UBC: Unipolar brush cells; ET: Essential tremor.

We used Scanpy^31^ and Harmony^32^ to integrate nuclei across donors, batches, sex and case-control status. Using previously known markers and mammalian single-cell atlases^33–35^, we annotated a total of 14 clusters representing all major cell clusters of the cerebellum (Figure 1D). All identified clusters expressed canonical markers of their respective cell-types: Granule cells (*RBOFX3*), Purkinje (*CALB1, ITPR1*), molecular layer interneurons (MLI) with two distinct populations (MLI_1; *PTPRK*, MLI_2, *NXPH1*), Purkinje layer interneurons (PLI; *NPXH2, SNTG1*), Golgi cells (*DLGAP2*), unipolar brush cells (UBC; *PTPRO, LMX1A*), Bergmann glia (*SCL1A3*), astrocytes (*GFAP*), microglia (*PLXDC2*), endocytes (*FLT1, ATP10A*), pericytes (*CEMIP, DLC1*), oligodendrocytes (*MBP*) and oligodendrocytes progenitor cells (OPC; *SOX6, PDGFRA*) (Figure 1D and 1F). As expected, granule cells were overrepresented in most samples as they represent around 50-70% of extracted cells in the cerebellar cortex (Figure 1E). Bergmann glia were the most common glial cells, followed by astrocytes. Proportion of Purkinje cells and interneurons remained somewhat stable across all samples (Figure 1E).

### Cell-type specific effects of cerebellar scQTLs

All 109 individuals subject to snRNA-seq were simultaneously genotyped. Following imputation, variant QC, and removal of non-European donors, 103 individuals and 5.3 million SNPs were kept for scQTL analysis (Supplementary figure 1). We restricted our analysis to SNPs with a minor allele frequency (MAF) greater than 0.05 and to those found within 1 megabase pair (Mb) of the transcription start site (TSS). We used *TensorQTL* ^36^ to map eQTLs in all of the 14 cell-types identified. Using both nominal and permutation mapping, we found a total of 3,120 eGenes, i.e. genes that were significantly regulated by at least 1 genetic variant (Figure 2A; Supplementary figure 2). A majority of eGenes were found in granule cells, the most abundant cell type in our dataset. As previously reported, the number of scQTLs per cell-type was correlated with the number of cells (Figure 2C). This is in line with the previous observation that increasing the number of sequenced cells could increase the discovery of scQTLs^2^. Most scQTLs were found upstream of the TSS, as expected, given their potential effects around transcription factor binding sites (Figure 2D). Similar distributions around the TSS were also previously observed in brain scQTLs^2^.

**Figure 2.**
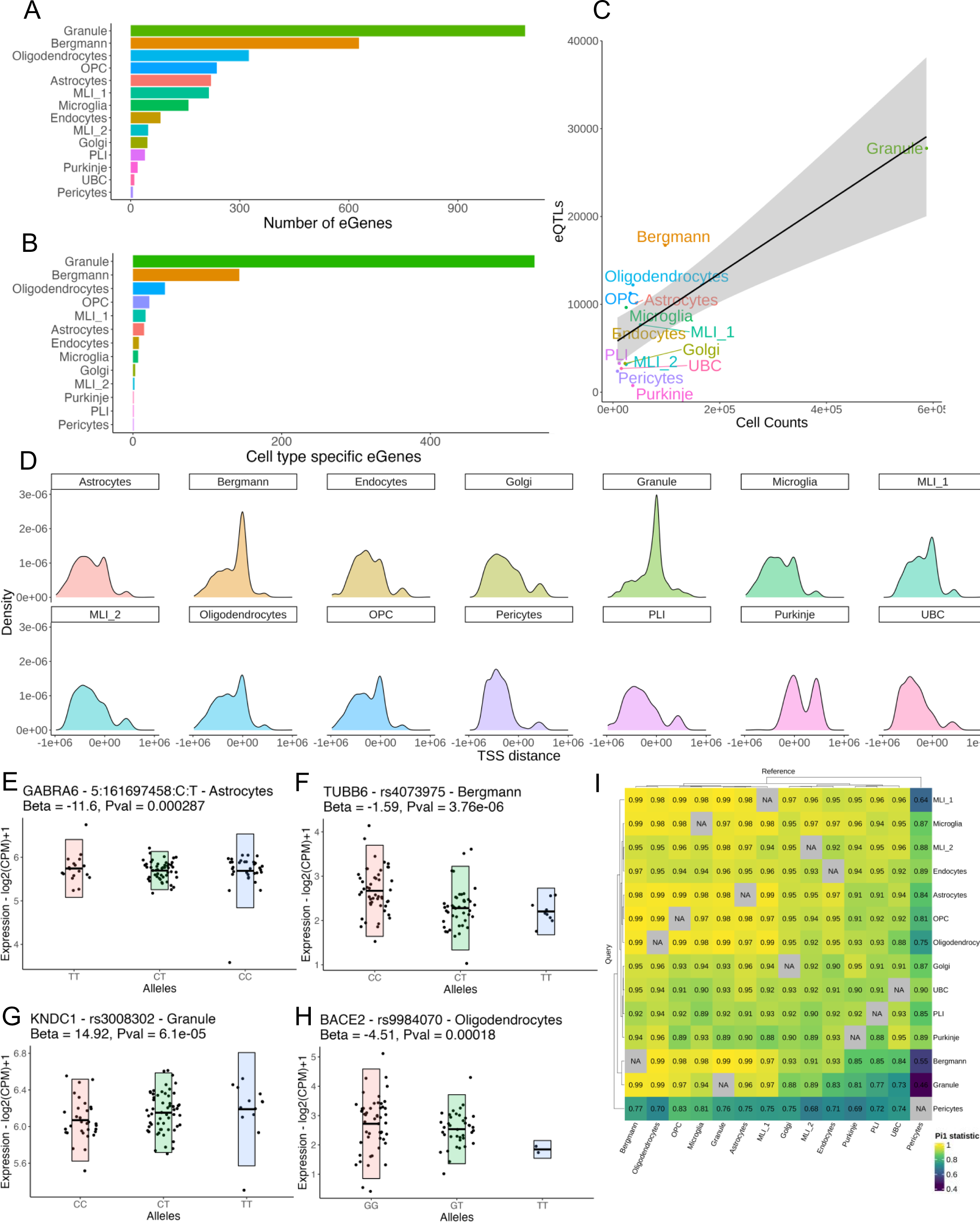
Identification of cell-type specific eQTLs in cerebellar nuclei. **A.** Number of eGenes per ce type. eGenes are defined as scQTL regulated genes passing FDR 5% using the permutation method *TensorQTL*. **B.** Number of cell-type specific eGenes across clusters. Cell-type specific eGenes are defined a genes regulated by scQTLs that pass FDR 5% in only 1/14 cell-types. **C.** Number of cell counts correlate wi number of discovered significant eQTLs (FDR 5%). **D.** Distribution of eQTLs in relation to the TSS. X-ax denotes distance from TSS (counted as 0 here) in base pairs. **E-H.** Boxplots of gene expression acros alleles for cell-type specific eGenes for Astrocytes, Bergmann glia, Granule cells and Oligodendrocytes. Eac point represents a different donor. Expression corresponds to log transformed counts per million (cpm). Blac line in boxplot corresponds to mean expression counts for a specific pair of alleles. Cell-type, top significa SNP for that eGene as well as Beta and p-values are indicated above the boxplot. **I.** Replication p-value across cell-types. Columns denotes reference cell-type whilst rows represent query cells for which Pi1 statist is calculated. Scale bar represents Pi1 statistic. **Legend:** MLI_1: Molecular layer interneuron 1; MLI_ Molecular layer interneuron 2; OPC: Oligodendrocyte progenitor cells; PLI: Purkinje layer interneuron; UB Unipolar brush cells; ET: Essential tremor.

We investigated shared effect sizes between cell-types in our dataset. For SNP-gene pairs at p-value < 1e-5, we studied the pairwise replication p-values for all cell-types (Figure 2I). Overall, cell-types shared a significant number of scQTLs between each other. (Pi1 0.73-0.99). Only pericytes, which had significantly lower number of scQTLs, had lower replication p-values (Pi1 0.46-0.89), possibly due to a small number of sequenced nuclei. Cell-types with the greatest number of scQTLs (and greatest number of cells) shared the most scQTLs with each other. For example, at p-value < 1e-5, granule cells displayed Pi1=0.99 with Bergmann glia. Glial cells shared the highest similarity with other glial cells. Interestingly, neuronal populations did not share higher Pi1 statistics with other neuronal populations. Of note, Purkinje scQTLs shared a large number of scQTLs with both UBCs and oligodendrocytes indicating some cross glial-neuron correlation between scQTLs. These results indicate that a majority of scQTLs are shared across different populations of cells and only a small fraction (<10%) represent cell-type specific eQTLs.

Next, we sought to study whether our single-cell results could replicate bulk-RNAseq eQTLs from the Cerebellar MetaBrain eQTL study^6^ (Supplementary figure 3). We used a similar approach as described above to calculate Pi1 statistic. Although most MetaBrain eQTLs were replicated in our dataset, scQTLs replicated at a lower rate when querying against MetaBrain eQTLs, possibly due to lower power in our analysis compared to the MetaBrain study. We find that granule cell scQTLs displayed the highest Pi1 statistic with MetaBrain eQTLs with over 50% of bulk eQTLs replicating in that population. Given the large number of granule cells in the cerebellum, this might indicate that most observed bulk eQTLs probably originate from these cells. Bergmann glia scQTLs replicated 41% of MetaBrain eQTLs, with other frequent cerebellar populations such as astrocytes, oligodendrocytes and OPCs replicating over 30% of bulk eQTLs.

We investigated whether our identified scQTLs had cell-type specific effects in the cerebellum. We defined cell-type specific eGenes as genes with significant genome-wide permutation tests in only 1/14 cell-types. We found a total of 803 cell-type specific eGenes in the cerebellum (Figure 2B). As with scQTLs, cell-type specific eGenes were correlated with the number of nuclei, with granule cells displaying over half of all cell-type specific effects. This further demonstrates the importance of large-scale sequencing in the discovery of scQTLs. We illustrate a few examples of cell-type specific eGenes (Figure 2E-H). For example, *KNDC1*, a guanine exchange factor known to regulate granule cell growth^37^, is specifically regulated in these cells. We also find that *BACE2,* a gene previously associated with ET and AD^24,38–42^, is specifically regulated in oligodendrocytes. *BACE2* was previously associated with oligodendrocyte- and endocyte-specific genetic regulation in an scQTL study of the human cerebral cortex^2^.

### Mendelian randomization of disease traits with cerebellar scQTLs identifies putative molecular mechanisms in neurological and psychiatric diseases

Disease associated SNPs often occur in non-coding regions and, therefore, understanding their molecular effects remains challenging. We used our newly constructed cerebellar scQTL dataset to investigate the causal effects between disease variants and expression events. We used summary-based mendelian randomization (SMR) to estimate the causal or pleiotropic relationship between GWAS SNPs and scQTLs. We also used the HEIDI test to distinguish pleiotropic effects from associations due to linkage disequilibrium (LD). We performed SMR for 5 traits: PD, AD, SCZ, Cerebellar volume (CBV) and ET (Figure 3A-D; 4B). We defined significant causal or pleiotropic associations as SNPs with FDR < 5% for the SMR test and SNPs with p-value > 0.05 for the HEIDI-test to eliminate LD associations.

**Figure 3.**
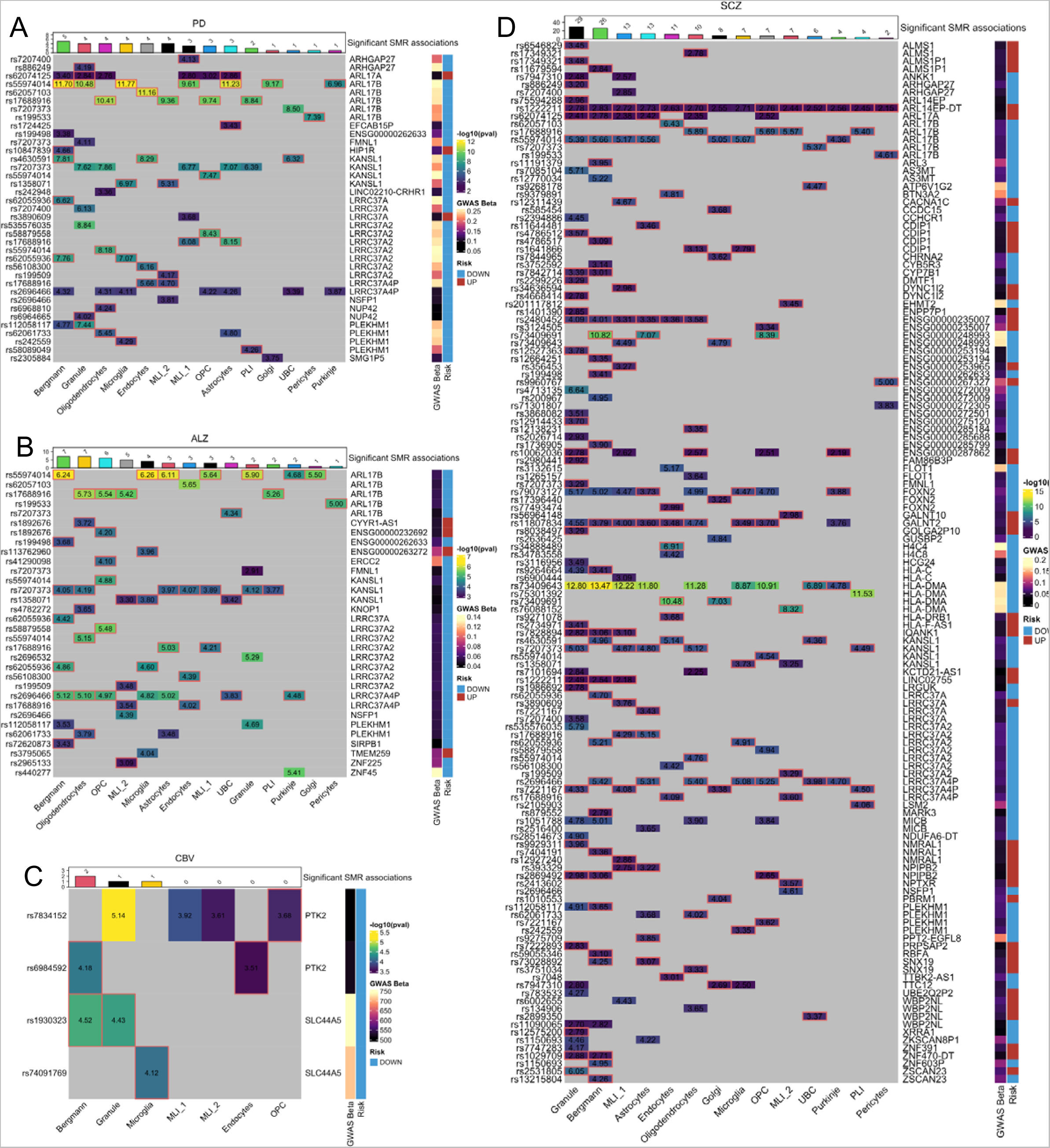
scQTL mendelian randomization for neurological and psychiatric disorders. **A-D.** Heatmap of summary-based mendelian randomization (SMR) results for A. Parkinson’s disease (PD), B. Alzheimer disease (AD), C. Cerebellar volume (CBV) and D. Schizophrenia (SCZ). Scale bar indicates SNP-gene pa SMR log10 transformed p-values. A highlighted red box around p-values indicates that SNP-gene pairs pas HEIDI test (p-value > 0.05). Rows indicate top SMR association SNPs. Columns indicate scQTL regulate genes. Row annotations indicate GWAS Beta size and risk direction (UP/red = increases disease ris DOWN/blue = decreases disease risk). Results are restricted to top SMR associations: PD, q-value = 0.0 AD, q-value = 0.05; CBV, q-value = 0.05; SCZ, q-value = 0.05. **Legend:** MLI_1: Molecular layer interneuron MLI_2: Molecular layer interneuron 2; OPC: Oligodendrocyte progenitor cells; PLI: Purkinje layer interneuro UBC: Unipolar brush cells.

**Figure 4.**
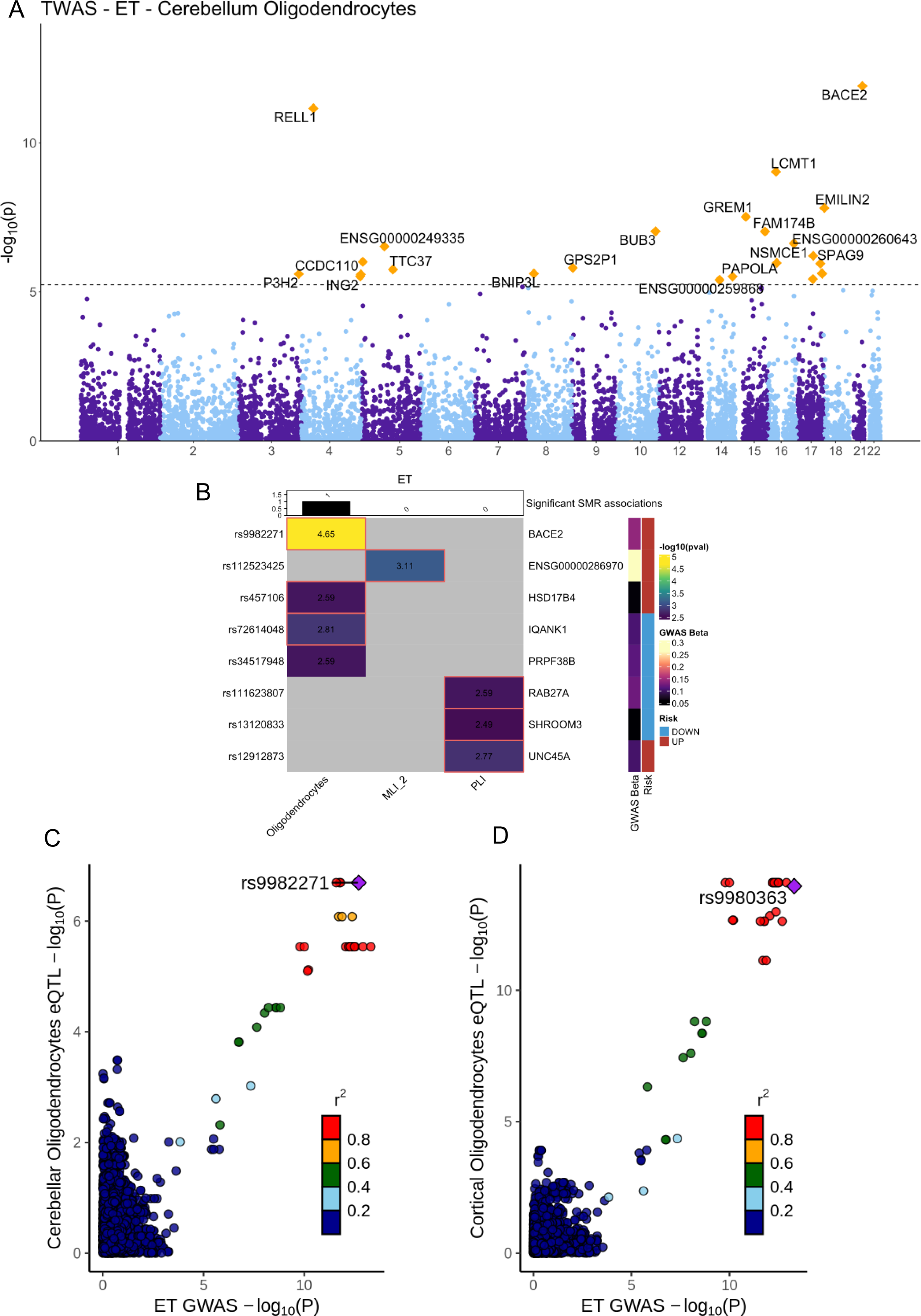
Oligodendrocyte-specific regulation of BACE2 expression in ET. **A.** Manhattan plot oligodendrocyte-specific TWAS for ET. P-values are obtained through ACAT-O meta-analysis across OTTER methods (P+T (0.001), P+T (0.05), lassosum, SDPR, and PRS-CS). Dotted line indicates 5% FDR threshol Orange dots indicate genes that pass FDR. **B.** Summary-based mendelian randomization results for E GWAS across cerebellar cells. Scale bar indicates SNP-gene pair SMR log10 transformed p-values. highlighted red box around p-values indicates that SNP-gene pairs pass HEIDI test (p-value > 0.05). Row indicate top SMR association SNPs. Columns indicate scQTL regulated genes. Row annotations indica GWAS Beta size and risk direction (UP/red = increases disease risk, DOWN/blue = decreases disease risk We restricted results to SNP-gene pairs with SMR association q-values > 0.5. **C-D.** Colocalization plots f cerebellar (C) and cortical (D) oligodendrocyte scQTLs and ET GWAS for the BACE2 locus. X-ax corresponds to ET GWAS log10 p-values and Y-axis corresponds to scQTL p-values in oligodendrocytes. D color corresponds to linkage-disequilibrium r^2^ values for correlated SNPs. Purple corresponds to top scQT SNPs in both cerebellar and cortical datasets. **Legend:** ET: Essential tremor.

For PD, a majority of SMR-causally associated scQTLs were found in the 17q21.31 locus, a region associated with multiple neurological disorders^43,44,45^. This locus includes genes such as *ARL17B*, *LRRC37*, *KANSL1*, *CRHR1* and *MAPT*. Such associations in this locus were observed in most cerebellar cell-types and were not cell-type specific, in concordance with previously observed colocalization in cortical scQTLs across glial and neuronal populations^2^. Previous association of *MAPT* regulation by astrocyte-specific scQTLs were not replicated in the cerebellum, correlating with the scarcity of tau accumulations in this brain region. Although we found few cell-type specific SMR associations for PD, we found significant association for Bergmann glia specific *HIP1R* expression with increased risk for PD. *HIP1R* encodes a regulator of clathrin-mediated endocytosis and a known interactor of Huntingtin ^46^.

In AD, we also found significant associations for multiple genes in the 17q21.31 locus. As with PD, associations within this locus were not cell-type specific. We, however, found that a majority of significant associations outside of this locus were cell-type specific. Notably, AD had multiple significant SMR associations in oligodendrocytes and OPCs. Additionally, we found significant association with cell-type specific *KNOP1* expression in oligodendrocytes, *ERCC2* in OPCs, *SIRBP1* in Bergmann glia and *ZNF225* in MLI_2. A majority of these SMR associations had variants with a decreasing effect on disease risk, consistent with previous findings of late cerebellar involvement in AD^19^. Moreover, we failed to find significant microglia-specific SMR associations, including *BIN1* colocalization, hinting at potential lack of AD risk factors in cerebellar microglia. Overall, for both AD and PD, a majority of causal SMR associations were correlated with decreased effects on disease risk, concordant with pathological observations of late cerebellar involvement in these disorders^47^ ^18,48^.

For CBV, we found 2 loci that had causal SMR associations with cerebellar scQTLs: *PTK2* and *SLC44A5* that were significant in granule cells and Bergmann glia, the most abundant cell-types in the cerebellum. *PTK2* encodes FAK, a cell adhesion tyrosine kinase implicated in synaptic branching known to be associated with cerebellar hypoplasia^49,50^.

SCZ was associated with a large number of granule and Bergmann glia eGenes. This is consistent with previous associations of SCZ with excitatory neurons^2,51^. SCZ risk genes such as *ANKK1*, *FLMN1* and *DMTF1* were all associated with granule-specific scQTLs. Interestingly, we found multiple interneuron-specific eGenes associated with SCZ: calcium-voltage channel *CACNA1C* in MLI_1, proton transporter *ATP6V1G2* in UBCs and the cholinergic receptor *CHRNA2* in Golgi cells.

### ET risk variants drive *BACE2* downregulation in oligodendrocytes

ET remains one of the most common neurological disorders, yet its pathophysiology is poorly understood. In our ET SMR analysis, we found that top ET risk variant rs9982271 was significantly associated with *BACE2* downregulation in oligodendrocytes (Figure 4B). We found that this scQTL was colocalized with ET risk SNPs at that locus (Figure 4C). Since ET genetic risk variants seem to be specifically enriched in the cerebellum, we tested whether scQTLs from cerebral oligodendrocytes also colocalized with ET. We found significant colocalization with rs9980363 and oligodendrocyte-specific scQTLs in the human cerebral cortex, hinting that *BACE2* regulation by ET risk variants might not be limited to the cerebellum (Figure 4D). To validate our SMR results, and further prioritize other ET risk genes, we performed cell-type specific transcriptome-wide association study (TWAS) using an Omnibus Transcriptome Test using Expression Reference Summary (OTTERS)^52^ (Figure 4A). We found that *BACE2* was prioritized as the most significant oligodendrocyte locus (ACAT-O p-value = 1.24e-12). Our results indicate that ET-associated variants in the upstream region of *BACE2* control its expression in cerebellar oligodendrocytes.

### Expression of ET-associated loci is significantly enriched in adult cerebellar oligodendrocytes and is independent from PD

Etiological mechanisms for ET pathophysiology are mostly unknown. Given this novel oligodendrocyte-ET association found through our scQTL analyses, we sought to validate these findings using our cohort of cerebellar snRNA-seq from ET patients. We first calculated the genetic enrichment for ET in our cerebellar dataset given that, other than *BACE2,* few GWAS loci colocalized with cerebellar scQTLs. We postulated that these loci might nonetheless display oligodendrocytes specific expression. We therefore used MAGMA cell-typing^51^ to calculate genetic enrichment within cerebellar cell-types. We found significant enrichment of ET loci in oligodendrocytes, Purkinje cells, PLIs and MLI_1 populations (Figure 5A). To control for correlated expression between cell-types, we ran MAGMA using conditional analyses to remove co-expression signal between cell-types. After conditioning on significant cell-types, we only found significant enrichment in oligodendrocytes. This enrichment was not replicated in fetal cerebellum or in adult cerebral cortex (Supplementary figure 4). These findings point to ET genetic risk driving disease progression specifically in adult cerebellar oligodendrocytes (Supplementary figure 4).

**Figure 5.**
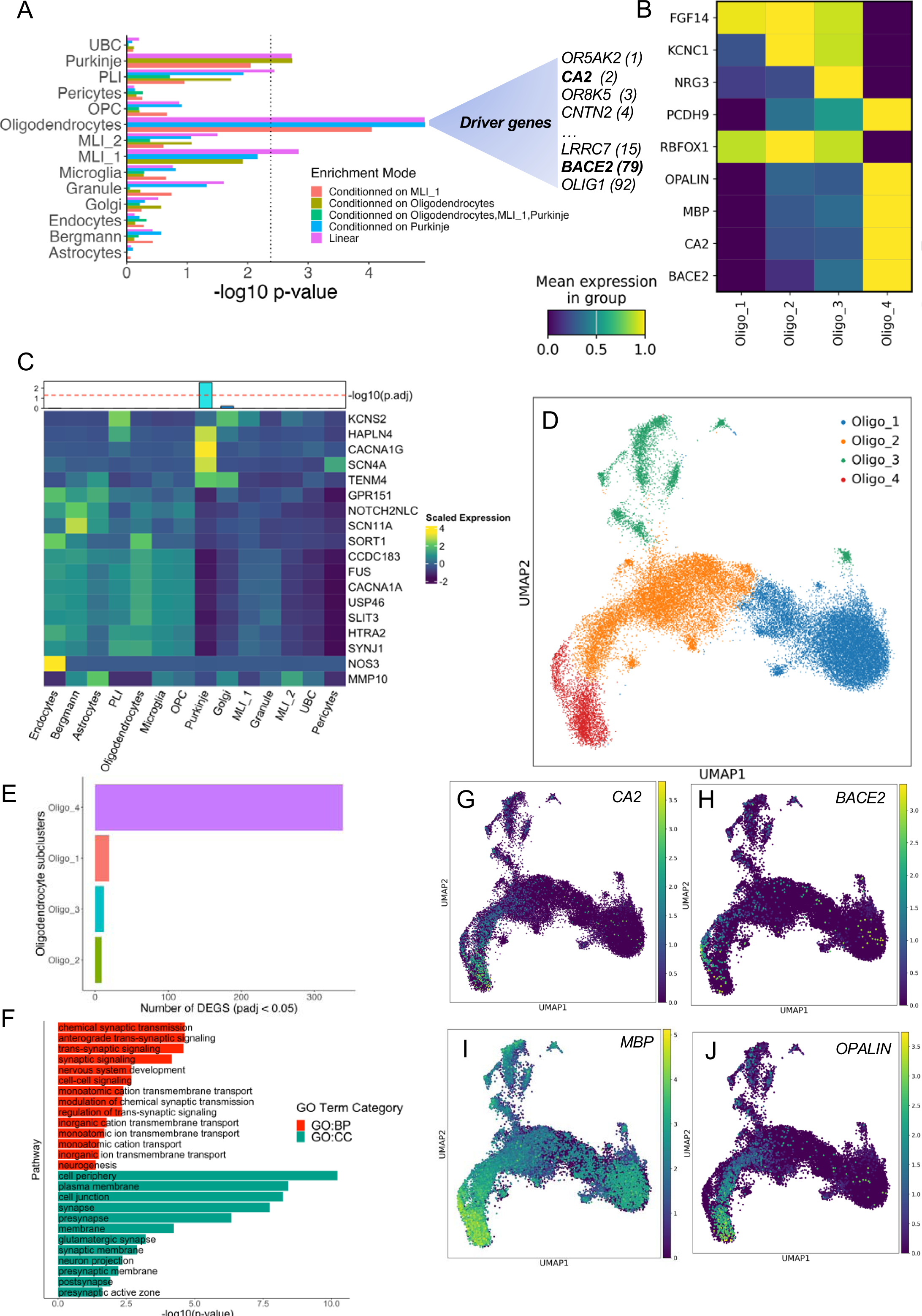
Genetic architecture of ET highlights vulnerability of oligodendrocytes and Purkinje cells. Bar plot of MAGMA genetic enrichment for ET associated loci in the cerebellar cortex. Colours indica calculation method: linear or conditioned for significant cell-types. Dotted line indicates 5% FDR. Driver gene are indicated on the right with rank in parentheses. **B.** ET associated loci from MAGMA and immatu oligodendrocyte markers are enriched in Oligo_4 subcluster. Scale colour represents mean normalize expression for each gene in each subcluster. **C.** Bar plot of log10 p-value and gene expression heatmap EWCE enrichment for genes harbouring rare variants in ET. Dotted red line in bagplot indicates 5% FD threshold. Heatmap depicts scaled expression for genes harbouring rare variants in ET across all 1 cerebellar cell-types. **D.** UMAP of oligodendrocytes subclusters. **E.** Bar plot of number of ET vs control DEG per oligodendrocyte subclusters. **F.** Pathway enrichment results for Oligo_4 DEGs. Colours denote Gen Ontology (GO) Term Category (BP: Biological pathway, CC: Cell communication). All terms pass 5% FD only top 12 terms per category are included. **G-J**. UMAP representation marker expression in Oligo_4. No the high expression of ET associated loci *CA2* and *BACE2* in this cluster as well as the high expression *OPALIN* and *MBP.* Scale bar represents normalized counts. **Legend:** MLI_1: Molecular layer interneuron MLI_2: Molecular layer interneuron 2; OPC: Oligodendrocyte progenitor cells; PLI: Purkinje layer interneuro UBC: Unipolar brush cells; ET: Essential tremor.

We thought this enrichment could be explained in part by genetic correlation between ET and PD. This is due to the fact that 1) PD loci were shown to be enriched in cortical and spinal oligodendrocytes^53^ and 2) ET and PD share a genetic correlation of 30%^24^. To test whether the enrichment of ET loci in oligodendrocytes is independent from PD, we conditioned the ET GWAS based on the most recent PD GWAS ^54^ to remove correlated signals (Supplementary figure 4). We found that ET loci were still significantly enriched in oligodendrocytes after conditioning on the PD GWAS. Furthermore, PD loci were not significantly enriched in cerebellar oligodendrocytes.

### Rare and common variants in ET are enriched in different cerebellar cell populations

We hypothesized that although ET common variants were enriched in oligodendrocytes, rare variants in ET might be enriched in other cell populations. We used an Expression Weighted Cell-type Enrichment (EWCE)^55^ test to perform gene-set enrichment analysis using a list of genes previously found to harbour rare variants associated with ET (*KCNS2, HAPLN4, CACNA1G, SCN4A, TENM4, GPR151, NOTCH2NLC, SCN11A, SORT1, CCDC183, FUS, CACNA1A, USP46, SLIT3, HTRA2, SYNJ1, NOS3, MMP10*). Interestingly, genes harbouring rare variants for ET were significantly enriched in Purkinje cells (Figure 5C). This could potentially explain the heterogeneity of clinical presentation in sporadic and familial cases of ET, with typical familial ET presents with early onset, compatible with a direct genetic effect on Purkinje cells^56^.

### A subset of immature oligodendrocytes drives genetic enrichment of ET loci

We hypothesised that a subset of cerebellar oligodendrocytes might display significant enrichment of ET risk genes. Using Harmony, we re-clustered cerebellar oligodendrocytes and identified 4 oligodendrocytes subclusters (Oligo_1; *FGF14*; Oligo_2: *KNC1*, Oligo_3: *NRG3*, Oligo_4: *PCDH9*) (Figure 5B & 5D). To study subtype enrichment, we first pulled the top 100 genes driving ET enrichment in oligodendrocytes. Surprisingly, *BACE2* was found at rank 79. Among the top 15 driving genes, we found oligodendrocyte markers such as *CA2, LRRC7* and *OLIG1*^57–59^(Figure 5B). These genes are concomitantly found in loci indicative of genome-wide significance on chromosomes 8, 1, and 21, respectively^24^. Most of these ET-associated genes, including *BACE2*, had enriched expression in Oligo_4 (Figure 5H). We also found that Oligo_4 displayed high expression of immature oligodendrocyte markers such as *OPALIN*, and low expression of mature markers such as *RBFOX1*^7^(Figure 5B & Figure 5I-J). Moreover, this cluster had high expression of *MBP*, indicative of active re-myelination (Figure 5I). Although both ET and controls had similar numbers of *RBFOX1+* Oligo_1 (adjusted p-value = 1.56e-1), we found that Oligo_4 cells were significantly more represented in ET samples than in control samples (permutation Z-score = 5.81, adjusted p-value = 2.57e-8; Supplementary figure 5).

We performed differential expression analysis across all 4 oligodendrocyte subclusters to identify ET driven changes in mRNA expression (Figure 5E & 5F). In line with ET loci enrichment in Oligo_4, this population displayed the strongest effect across the 4 subtypes (Oligo_4: 339 DEGs; Oligo_1: 19 DEGs; Oligo_3: 12 DEGs; Oligo_2: 9 DEGs) (Figure 5C). We performed pathway enrichment analysis of Oligo_4 specific DEGs and found significant enrichment of pathways related to synaptic signalling and ion transport across neuronal membranes (Figure 5F). These processes are known to be crucial to myelin and axonal homeostasis ^60,61^ ^62^ and could be important compensatory mechanisms in ET oligodendrocytes. Overall, these results indicate that a subset of genetically vulnerable oligodendrocytes display dysfunctional synaptic and ionic changes in ET.

### Neuronal dysfunction and Bergmann gliosis in the ET cerebellar cortex

Oligodendrocyte vulnerability alone is unlikely to drive disease progression in ET. We therefore sought to assess other affected neuronal and glial populations in the cerebellar cortex. Using pseudobulk differential expression, we observed the strongest effects in glial cells, mostly Bergmann glia, OPC, and astrocytes (Figure 6A). Concordant with the finding of Bergmann gliosis in the ET cerebellar cortex ^63^ ^64^ ^65^, Bergmann glia displayed the largest number of DEGs. Bergmann DEGs were mostly related to gliogenesis and axon projection (Figure 6D). Multiple genes related to gliosis such as *ID4, GFAP, VIM* and *NES*^66^, were all found to be significantly upregulated in Bergmann glia (adjusted pval < 0.05, log2FC > 0.5) (Figure 6C). Given the lack of genetic enrichment of both common and rare variants in this population, we postulated that the effects observed in this population are secondary to neuronal insult as is observed in spinocerebellar ataxia^67^.

**Figure 6.**
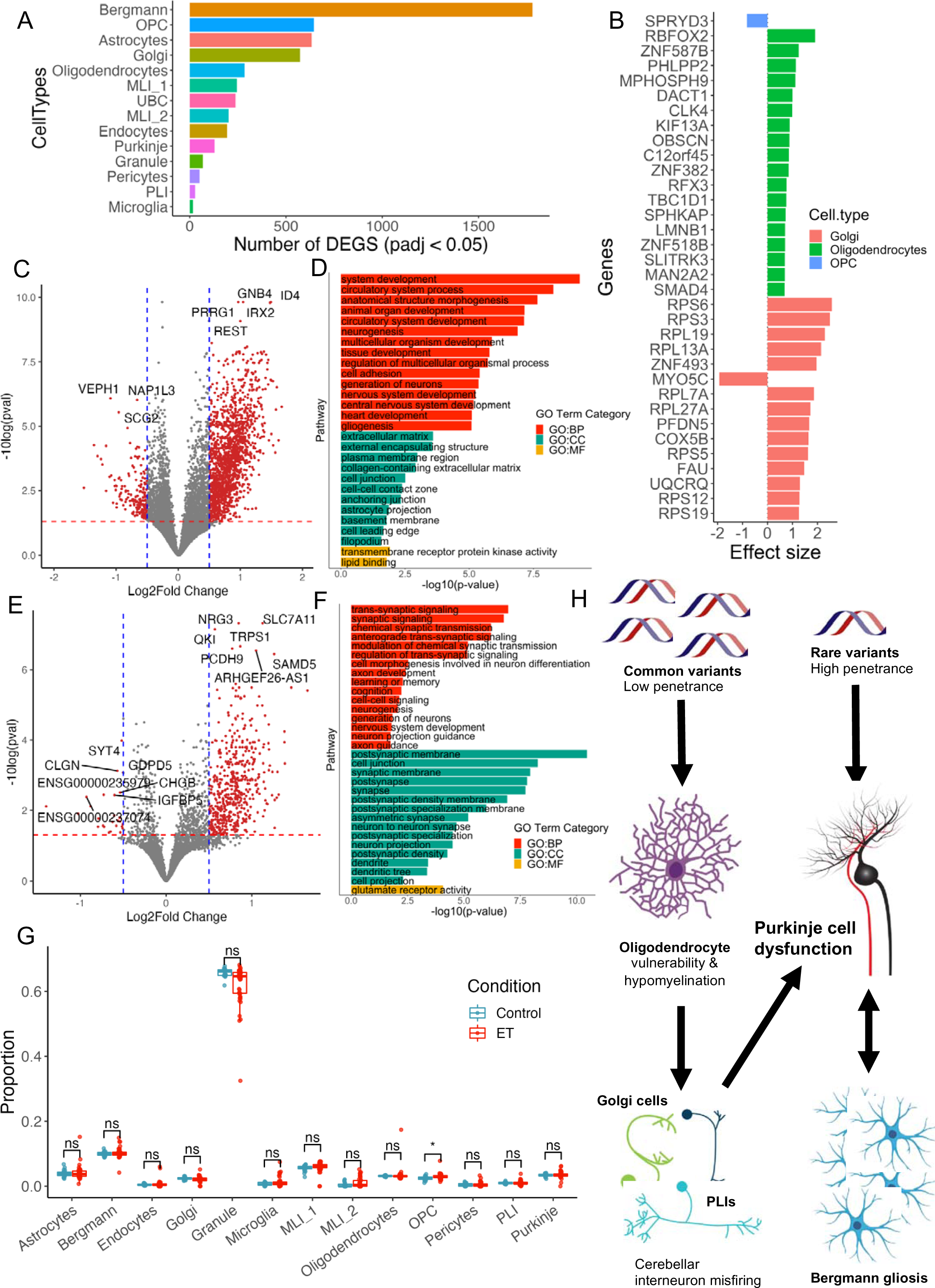
Differential expression highlights oligodendrocyte and Golgi cell dysfunction in the E cerebellum across multiple cohorts. **A.** Bar plot of number of ET vs control DEGs per cell-type for cerebellar clusters. Strongest effects are observed in glial populations and in Golgi neurons. **B.** Bar plot cell-type DEGs from bulk-RNA sequencing of 33 ET and 21 control samples. All genes pass FDR (25%). axis indicates effect size and direction. Top 18 genes from each cell-type are included, irrespective of effe direction. **C and E.** Volcano plot of Bergmann glia DEGs (C) and Golgi DEGs (E). X-axis indicated log fo change and Y-axis indicates –log10 p-value. Red line indicates 5% FDR. Blue line corresponds to -0.5 an 0.5 log2 fold change. **D and F.** Pathway enrichment analysis for Bergmann glia DEGs (D) and Golgi DEG (F). Colours denote Gene Ontology (GO) Term Category (BP: Biological pathway, CC: Cell communicatio MF: Molecular function). All terms pass FDR 5%. Top 15 significant terms per category are included. Boxplot of differences between cell-type proportions for ET and control bulk-RNA seq samples (n = 49 ET n= 37 controls). Note the small but significant increased proportion of OPCs in ET vs controls (adjusted value = 0.032). Boxplot center line represents median of group, upper and lower limits represent first and thi quartiles of data. Whiskers represent interquartile range x1.5. **H.** Postulated pathophysiological mechanism play in the ET cerebellar cortex based on current snRNA-seq results. **Legend:** MLI_1: Molecular lay interneuron 1; MLI_2: Molecular layer interneuron 2; OPC: Oligodendrocyte progenitor cells; PLI: Purkin layer interneuron; UBC: Unipolar brush cells; ET: Essential tremor.

Bergmann gliosis in ET is thought to arise around sites of Purkinje cells degeneration or loss. Interestingly, we found few DEGs in Purkinje cells when comparing ET to control samples (Figure 6A) despite sequencing a large number of Purkinje nuclei (37087 total Purkinje cells; 31,241 control Purkinje cells; 5,846 ET Purkinje cells (15.76%)). DEGs were mostly related to the post-synaptic membrane (adjusted p-value = 8.20e-03; Supplementary figure 6) including genes such as *LRRTM4, GLRA3* and *TENM2*. This is in line with previous transcriptomic characterization of ET Purkinje cells, where only 36 genes were found to be dysregulated compared to healthy Purkinje cells^68^. Golgi cells were found to be the most affected neuronal population in our cohort (Figure 6A; 575 DEGs with adjusted p-value < 0.05 & log2FC > or < 0.5). Golgi cells are the most abundant inhibitory interneurons and are crucial for the inhibition of granule cells^69^. We found that Golgi cell DEGs were related to synaptic transmission, axonal development, dendritic tree, and glutamate receptor activity (Figure 6E & 6F). Given these findings we sought to investigate the differential cell-cell connections between Golgi cells and other cerebellar neurons using CellPhoneDB^70^. We found that ET Golgi cells had differential cell-cell communications with most cerebellar neurons, including granule cells, mainly through Teneurin- and Ephrin-signalling proteins (Supplementary figure 7). We found the strongest effect in ET differential cell communications to be between Golgi cells and PLIs via *TENM2/3/4-ADGRL2* adhesions. Concomitantly, studying the cell-cell communications between oligodendrocytes subclusters and neurons highlighted numerus specific Oligo_4 and PLI communications through leucin-rich repeat (LRR) proteins and neurexins. These transmembrane proteins are known to be involved in myelination and axonal differentiation. In-depth functional characterization of these populations in both the healthy and ET cerebellum are lacking, but our results indicate that Golgi cells might display dysregulated synaptic communications with PLIs in ET.

### Oligodendrocytes and Golgi cells are dysregulated in an independent ET cohort

Our single cell sequencing of ET patients is limited by sample size (n = 16). We therefore sought to validate our DGE findings in an independent cohort of ET cerebellar cortex. Given observed loss of Purkinje cells in histopathological studies of ET, we were interested in: 1) evaluating consistent changes in cell-type proportions across cohorts and 2) correlate our cell-type specific DEGs across snRNA-seq and bulk RNA-seq. We deconvoluted publicly available bulk-RNA seq from 33 ET cases and 21 controls (Columbia cohort, <u>GSE134878)</u> using Bisque^71^ to estimate cell-type proportions. We also deconvoluted bulk-RNA reads from the same 16 ET and 16 control cerebellar samples described in this study (Saskatchewan cohort). Using permutation testing and a meta-analysis approach across these 2 ET cohorts, we tested for cell-type proportion differences between case and control samples (Figure 6G). Interestingly, we found no differences in the proportion of Purkinje cells between ET and control samples. However, we found an increased proportion of OPCs (Figure 6G; Stouffer’s Z = 3.03, adjusted p-value = 0.032) in ET compared to controls across both cohorts, in line with our current observations of vulnerable oligodendroglial cells. We used the deconvoluted bulk-RNA reads from the Columbia cohort to estimate the cell-type specific effect of bulk-RNA seq DGE results using BSEQ-SC^72^ (Figure 6B). Deconvoluted DEGs were uniquely discovered in Golgi, oligodendrocytes and OPCs. We found multiple genes related to oligodendrocyte differentiation and myelination such as *RBFOX2* and *LMNB1*^73^. Two genes from our snRNA-seq DGE analysis, *FMRD4A* and *CCDC30*, were also replicated in our deconvolution analysis for Golgi cells with same-direction LFC (BSEQ-SC FDR 25% & adjusted p-value < 0.05). Our replication of oligodendrocyte and Golgi dysfunction in independent ET cohorts increases confidence in our initial findings on the vulnerability of these cell populations.

## Discussion

Most disease-associated loci have cell-type specific effects and thus characterizing these molecular mechanisms is crucial to understanding disease pathophysiology and for therapeutic development. Brain diseases are subject to both cell-type and region-specific genetic effects. The mapping of scQTLs in different brain regions is therefore important to identify causal cell-types in neurological disorders. In this study, we used large-scale snRNA-seq coupled with genotyping to build an scQTL atlas of the human cerebellum. Although previous efforts of scQTL mapping in the brain had leveraged multiple cortical regions^2^, we limited region-specific effects on gene expression by studying gene regulation in a single region. This limits possible underlying biases caused by region-driven gene expression as was previously shown.

We found that cerebellar scQTLs shared multiple similarities with cortical scQTLs. The number of scQTLs was correlated with the number of sequenced cells. As such, given their high cellular density in the cerebellum, granule cells displayed the largest number of scQTLs. Unsurprisingly, we found that granule cell scQTLs accounted for more than half of bulk RNA-seq eQTLs, suggesting that a majority of eQTLs from whole transcriptomic studies miss eQTLs in rarer cell types in the cerebellum. Even through the use of snRNA-seq, a majority (50-70%) of sequenced nuclei are granule cells. This limits the discovery of scQTLs in rarer cell populations given that mapping is highly dependent on number of sequenced cells. While this could explain the lower number of scQTLs in Purkinje cells and interneuronal populations of the cerebellum, these observations cannot be attributed to just statistical power limitations. This is evidenced through the fact that for an equivalent number of sequenced cells, glial cell populations displayed more scQTLs than neuronal populations (with the exception of granule cells). It could be argued that expression in cerebellar neurons, especially Purkinje cells, might be more “immune” to genetic regulation, given their important role in motor coordination. Genetic variation might have a lesser effect on gene expression in these cells than in others from glial cell lineages.

Non-coding genetic variants are the most common disease-associated risk factors in neurological disorders. Understanding how and where these variants mediate their effects remains a major subject of investigation in disease genomics. Here, we investigated causal relationships between scQTLs and GWAS associated variants for multiple neurological and psychiatric traits including ET, the most common cerebellar disorder. We replicated known disease-cell associations such as oligodendrocytes with AD^74^ and excitatory neurons with SCZ^75,76^. In the latter, glutamatergic granule cells could contribute to known dysfunctional connections between the cerebellum and the dorsolateral prefrontral cortex^75,77^. For AD and PD, most causal SNP-gene associations were negatively correlated with disease risk, hinting at potential protective effects of cerebellar scQTLs. This could correlate with the relatively spared cerebellum in these disorders^18,47^.

Causal cell-types in ET remains a subject of controversy. Previous histopathological studies of the ET cerebellum have identified selective Purkinje cell degeneration and loss, although this finding remains disputed^29,78^. Furthermore, whether this process is cell type autonomous or implicates dysfunction of other cerebellar cells is not known. Using our scQTL atlas, we found a possible causal association between ET common variants and a population of immature actively myelinating oligodendrocytes. This population was shown to be more frequently found in ET brains and have decreased expression of ion channels associated with myelin and axonal homeostasis. Oligo_4 was also found to mostly interact with cerebellar interneurons such as PLIs, hinting that that these populations might suffer from deficient myelination. These findings correlate with known demyelination of the cerebellar peduncles in ET^79–82^, a feature that differentiates this disorder from PD^83^. Moreover, ET associated variants in the *BACE2* locus are also associated with radial diffusivity abnormalities in the inferior cerebellar peduncles^24^. Although both AD and PD have been associated with oligodendrocyte dysfunction, the cerebellar specific enrichment of ET variants could explain phenotypical differences between these disorders. Concomitantly, lesions of cerebellar white matter are known to be associated with postural tremor, such as in multiple sclerosis^84^ and central hypomyelination syndromes such as leukodystrophies^85^. Our analyses also highlighted the possible genetic effects of ET rare variants on Purkinje cells. The direct effects of these variants on Purkinje functionality could explain the clinical differences between familial and sporadic forms of ET in terms of disease onset and progression^56^. Whether the vulnerability of cerebellar oligodendrocytes contribute to Purkinje cell dysfunction directly, or through other cerebellar interneurons (such as Golgi cells) is still unknown. Based on our current results, we formulated a putative disease model for ET in which rare and common variants converge on the dysfunction of Purkinje cell towards tremor generation (Figure 6H). Spatial characterization of interactions between vulnerable oligodendrocytes and cerebellar neurons could be required to understand the pathological processes underlying tremor in ET.

Our study has several limitations. Although our overall number of sequenced nuclei is large, the number of donors is smaller than other scQTL studies done in brain or in blood cells. As previously mentioned, Granule cells comprise a majority of sequenced nuclei and many neuronal and glial cell populations in our dataset are limited in sample size (< 50K cells). Although we identified *BACE2* driven mechanisms in ET, other ET GWAS loci failed to colocalized with cerebellar scQTLs, most notably the only replicated SNP across two GWAS studies, rs17590046, found in the region of *PPARGC1A* on chromosome 4. Given our limited sample size, a number of cerebellar scQTLs may remain undefined. Moreover, the cerebellum is host to numerous splicing events which have been shown to be linked to synaptic regulation^86^. It could be argued that these ET variants may act through splicing QTLs to confer increased disease risk. Such studies could further our understanding of cerebellar function in both health and disease.

Our study highlights the importance of studying region-specific and cell-type specific eQTL events. Such information can reveal hidden disease-driving mechanisms underlying brain disorders. We show the utility of such analyses through our identification of a vulnerable oligodendrocyte population associated with ET in the human cerebellar cortex. We anticipate that further sub-regional characterization of brain cell scQTLs will be required to fully understand how genetic risk factors act on neuronal networks in the brain. We also hope that our dataset will be useful for future studies on cerebellar populations.

## Methods

### Sample collection

Frozen post-mortem cerebellar cortex tissue samples from 109 patients were obtained from the Movement Disorder Clinic Saskatchewan and the Douglas-Bell Brain Bank. Our ET cohort was previously described^87^. All patients underwent post-mortem neuropathological examination to validate clinical presentation. We selected healthy patients with no signs of neurological or psychiatric diseases. ET patient had a definite or probable diagnosis before death. Both ET and control samples were assessed based on the following criteria: unremarkable cerebellum, no cerebellar degeneration, no signs of Parkinson’s disease or other movement disorders, mild signs of Alzheimer’s disease.

### Nuclei extraction, barcoding, and sequencing

Nuclei from frozen tissues were extracted using a modified version of the SPLiT-seq protocol^30^. Briefly, samples were homogenized in Nuclei Isolation Media (containing 0.4 U/uL Enzymatic RNAse-In and 0.2 U/uL SUPERase RNAse inhibitors) using Dounce homogenizers. Lysates were filtered through 40-um cell filters and centrifuged at 400 rpm for 5 minutes. After resuspension in 1X PBS, samples were filtered once again in 40-um filters before fixation using a 1.33% Formalin solution. Nuclei are then permeabilized using 5% Triton solution. Cells were then counted using a hemocytometer before barcoding. Barcoding and sequencing were performed at the Johns Hopkins Sequencing Core. Nuclei cDNA was barcoded using the Evercode WT and Mega kits from Parse Bioscience. Samples were sequenced in 3 different batches (WT, Mega_01 and Mega_02). WT batch (which was used for optimization) was comprised of 80,000 nuclei. Each Mega batches were comprised of 500,000 nuclei with 64 and 45 samples respectively. Nuclei were sequenced on the NovaSeq 6000 S2 (WT) and S4 (Mega_01 and Mega_02) flow cells aiming to obtain at least 20K reads per cell.

### Single-nuclei RNA-seq analysis

Reads from all batches were processed using the split-pipe software v0.9 (default settings with min_cells set at 30K) using the Ensembl Homo_sapiens. GRCh38.105 as reference annotation. All batches were merged and analyzed using Scanpy^31^. Nuclei with fewer than 450 UMIs, over 10K features and 10% mitochondrial reads were removed. Nuclei were integrated across donors, extraction batches, sequencing batches, case/control status and sex using Harmony^32^ (default settings with max iterations set at 20). Nearest neighbor analysis was performed using the cosine metric. Following UMAP and Leiden clustering (resolution = 1), cluster annotation was performed using known marker genes base on differential gene expression using the Wilcoxon test. This resulted in the annotation of 14 major cell-types of the cerebellum. We performed subcluster analysis for cerebellar oligodendrocytes using Scanpy, selecting all oligodendrocytes from the original 1M cell object. We integrated across batches, case/control status and sex using Harmony (iterations = 20) and clustered using Leiden clustering (resolution = 0.1) identifying 4 distinct clusters. Cluster markers were determined using the Wilcoxon test as implemented in scanpy.

### Genotyping

DNA was extracted from 109 cerebellar cortical samples using the Qiagen DNeasy kit. using the Global Screen Array GSA 3.0 from Illumina was used to genotype all samples. Data was loaded into GenomeStudio and standard quality control (QC) procedures were applied to remove poor quality data ^88^. Briefly, variant level QC removed variants with the following: genotyping call rate of < 95%, minor allele frequency (MAF) < 0.01, and variants failing Hardy-Weinberg equilibrium (10e-6 threshold). Samples with individual call rates of < 98%, discordant gender, failing heterozygosity tests, were related to other samples (PI_HAT > 0.2), or were of non-European ancestry were removed. Ancestry was inferred through principal component analysis (PCA) where European samples clustered with European sample from 1000 genome reference panel (Supplementary Figure 1). This left 103 samples covering 467,477 variants passing quality control. Imputation was then done using the Michigan Imputation Server^89^ using the Eagle v2.4^90^ and the Haplotype Reference Consortium Reference Panel version 1.1^91^.

### scQTL mapping

We used TensorQTL^36^ to map eQTLs in all 14 cell-types in our dataset. We first generated pseudobulk RNA-seq profiles for all 14 clusters using DESeq2 ^92^ by aggregating counts for each gene across each cell-type. Counts were aggregated across all batches for each given individual. Counts were normalized using TMM as implemented in EdgeR^93^. We restricted our analysis to genes with an average expression of at least 1 count per million. We also limited our analyses to SNPs with MAF > 0.05. For all scQTL mapping methods we used the following covariates: first 3 genotyping PCs, age, sex, case/control status and the first 30 expression PCs. Phenotype-level summary statistics were obtained using permuations (cis.map_cis). All variant-phenotype associations were assessed using the nominal mapping mode (cis.map_nominal). P-value histograms were assessed for uniform distribution for all cell-types.

### Pi1 statistic

To assess SNP-gene pair p-value replication between scQTLs and cerebellar MetaBrain eQTLs, we calculated the pi1 statistic between both datasets using the q-value package. We selected SNP-Gene pairs indicative of significance with p-value < 1e-5 and selected corresponding p-values for the same SNP in the query cohort. We used both SNP-gene pairs from the MetaBrain study and cerebellar scQTLs as reference and query. We also used a similar approach to assess replication across cell-types, using each cell populations as either reference or query.

### Summary-based Mendelian Randomization

To assess the effects of scQTLs on disease risk, we used the SMR and HEIDI tests as available in the SMR software tool^94^. SMR implements mendelian randomization methods to test whether the effects of a disease-associated variant are mediated through gene expression changes. For all cell-types, we tested for associations between significant GWAS SNPs for various traits (ET, PD, AD, SCZ, CBV) and scQTLs (with p-value < 1e-05). GWAS summary statistics for ET were obtained through 23andMe, Inc. Summary statistics for PD ( <u>GCST009325</u>) ^54^, AD (<u>GCST90027158</u>) ^95^ and CBV ( <u>GCST90105075</u>) ^96^ were obtained through the GWAS catalog. SCZ GWAS summary statistics were obtained from the Psychiatric Genetics Consortium ^97^. We differentiated causal or pleiotropic associations from those due to linkage using the HEIDI test (p-value > 0.05) using an LD-reference from the 1000 Genomes Project (restricted to individuals of European ancestry).

### Cell-type specific TWAS

We used OTTERS^52^ to perform cell-type specific TWAS. OTTERS uses polygenic-risk score methodology to 1) create expression weights based on summary eQTL data and 2) perform an omnibus test which studies the effects of SNPs on gene-expression. It has been shown to be more powerful for TWAS as other methods using individual-level data. For each cell-type, we generated SNP-level expression weights using cerebellar scQTLs and an LD reference data from the 1000 Genomes Project. We generated weights using 5 different models: P+T (0.001), P+T (0.05), lassosum, SDPR, and PRS-CS. We then performed summary-level TWAS for ET using GWAS summary statistics and cell-type specific expression weights. Using phenotype-level TWAS results across all 5 models, we performed an ACAT-O ^98^ test to aggregate p-values across all models.

### MAGMA cell-typing and EWCE rare variant enrichment

We calculated the enrichment for ET heritability in our cerebellar snRNA-seq dataset using the *MAGMA* R package. We first processed ET GWAS summary statistic using *MungeSumstats* to then build a gene-level signature for cell-type enrichment. After converting our cerebellar Seurat object to CTD, we used both linear and conditional (conditioning on all significant cell-types) to calculate cell-type enrichment in our cerebellar cell-types. We also calculated cerebellar enrichment for PD GWAS (‘ieu-b-7’) using both conditional and linear enrichment. We then re-calculated ET GWAS enrichment after controlling for the PD GWAS. Finally, we calculated ET GWAS enrichment in both fetal cerebellum (<u>GSE165657</u>)^33^ and cortical M1 single-cell data from the Allen Brain Institute (https://portal.brain-map.org/atlases-and-data/rnaseq/human-m1-10x). We also used the EWCE^55^ R package to calculate the enrichment of genes harbouring rare variants in ET. We built a list of genes from previously published familial, rare variants and linkage studies in ET^99^. We calculated the cell-type enrichment for those genes using a bootstrap enrichment test using 10000 random lists as implemented in the package.

### Pseudo-bulk differential expression analysis

We used DESeq2^92^ to perform pseudobulk differential expression. We extracted mRNA counts per cell-type and sample to construct pseudobulk count matrices each cell population. We aggregated counts across different sequencing batches for the same sample. We removed genes expressed in less than 10 cells. Batch effects were assessed using PCA for each cell-type. We used the Wald-test using condition, age, and sex as covariates to test for DEGs across case-control status for all cell-types. For oligodendrocyte subcluster differential expressed, we found significant sequencing batch effects across subclusters and added sequencing batch as a covariate in the Wald-test. All DEGs were controlled for 5% FDR.

### Cell-cell communication analysis

We used CellPhoneDB^70^ to predict differential cell-cell communications between ET and control cells in the cerebellum. To map differential cell-cell interactions between ET and controls, we used the DEG-based method, using upregulated significant DEGs (adjusted p-value < 0.05 & Log2FC > 0) for all 14 cell-types. We also mapped wild-type cell-cell communications between oligodendrocyte subclusters (Oligo_1, 2, 3 and 4) and cerebellar neurons using the statistical analysis method.

### Deconvolution of ET bulk-RNA sequencing

FASTQ files from bulk RNA-sequencing of 33 ET samples and 21 controls (GEO accession number: <u>GSE134878</u>; Columbia cohort) were mapped to Ensembl Homo_sapiens.GRCh38.105 annotation of the human genome using Salmon v1.4.0^100^. We also mapped FASTQ files from anther cohort of 16 ET patients and 16 controls subject to snRNA-seq in this study (Saskatchewan cohort). We used Bisque^71^ to deconvolute reads from all 70 samples (49 ET and 37 controls) which allows to estimate the proportion of cell-types within whole transcriptome samples. We compared cell-type proportions using permutation testing of linear regressions using age, sex, and case/control status as covariates (10K permutations without replacement). We used Stouffer’s Z method^101^ to combine Z-scores across both cohorts (Columbia cohort and Saskatchewan cohort). P-values were corrected for FDR 5%. For the Columbia cohort, we used cell-type proportions to estimate cell-type specific effects of DEGs between ET and control samples. Briefly, we first used DESeq2 to perform DGE analysis between ET and control tissues using age, sex, and case/control status as covariates. We also performed DGE analysis using a model including the following cell-type proportion as covariates: Purkinje, Bergmann glia, Oligodendrocytes, Astrocytes, OPC and Golgi. Cell-types included in this model were selected based on effect sizes in snRNA-seq pseudobulk DGE analyses. We selected genes that: 1) displayed adjusted p-values < 0.05 in both analyses and 2) displayed more significant adjusted p-values when including cell-type proportion as covariates than in the base model. For each of these genes, we used BSEQ-SC to test whether their regulation was cell-type specific controlling for FDR 25%.

## Acknowledgements

Funding & Acknowledgments

We thank the Douglas Bell Canada Brain Bank at McGill for the acquisition of post-mortem cerebellar samples. We would also like to thank the research participants and employees of 23andMe, Inc. for making this work possible. CEC is supported by graduate scholarships from the Canadian Institutes of Health Research (CIHR) and the Fonds de Recherche du Québec en Santé (FRQS). MM is supported by a graduate scholarship from FRQS. ZS received a doctoral student fellowship from the CIHR Frederick Banting & Charles Best Canada Graduate Scholarship (FRN260055) and the Transforming Autism Care Consortium, a thematic network supported by the FRQS. This work is funded by grants obtained from the International Essential Tremor Foundation (IETF) and CIHR Project Grant.

