## Supplementary figures 1-7 for "A single-cell eQTL atlas of the human cerebellum reveals vulnerability of oligodendrocytes in essential tremor"

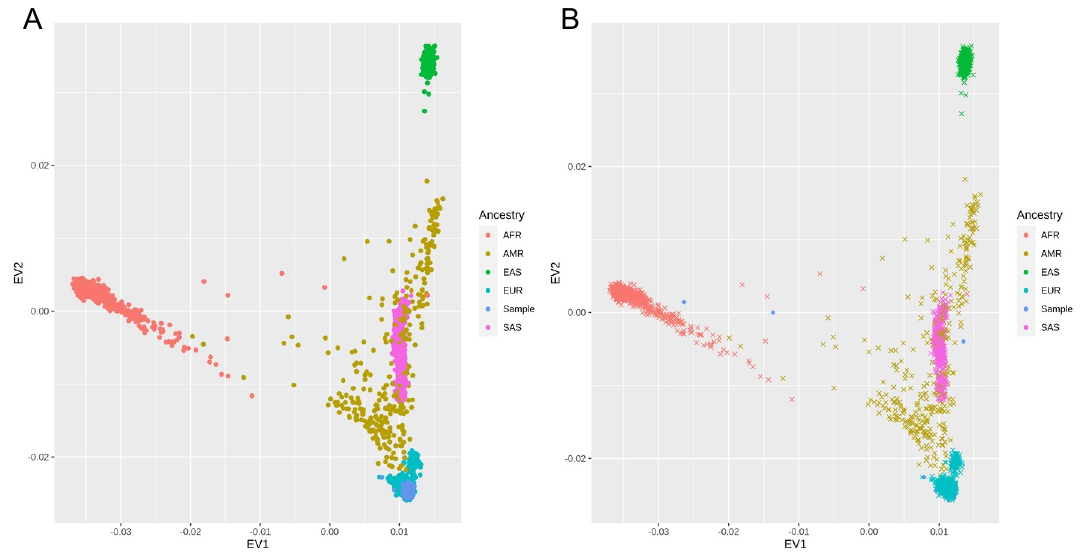

**Supplementary figure 1.** Ancestry PCA. Before (A) and after (B) removal of non-European donors.

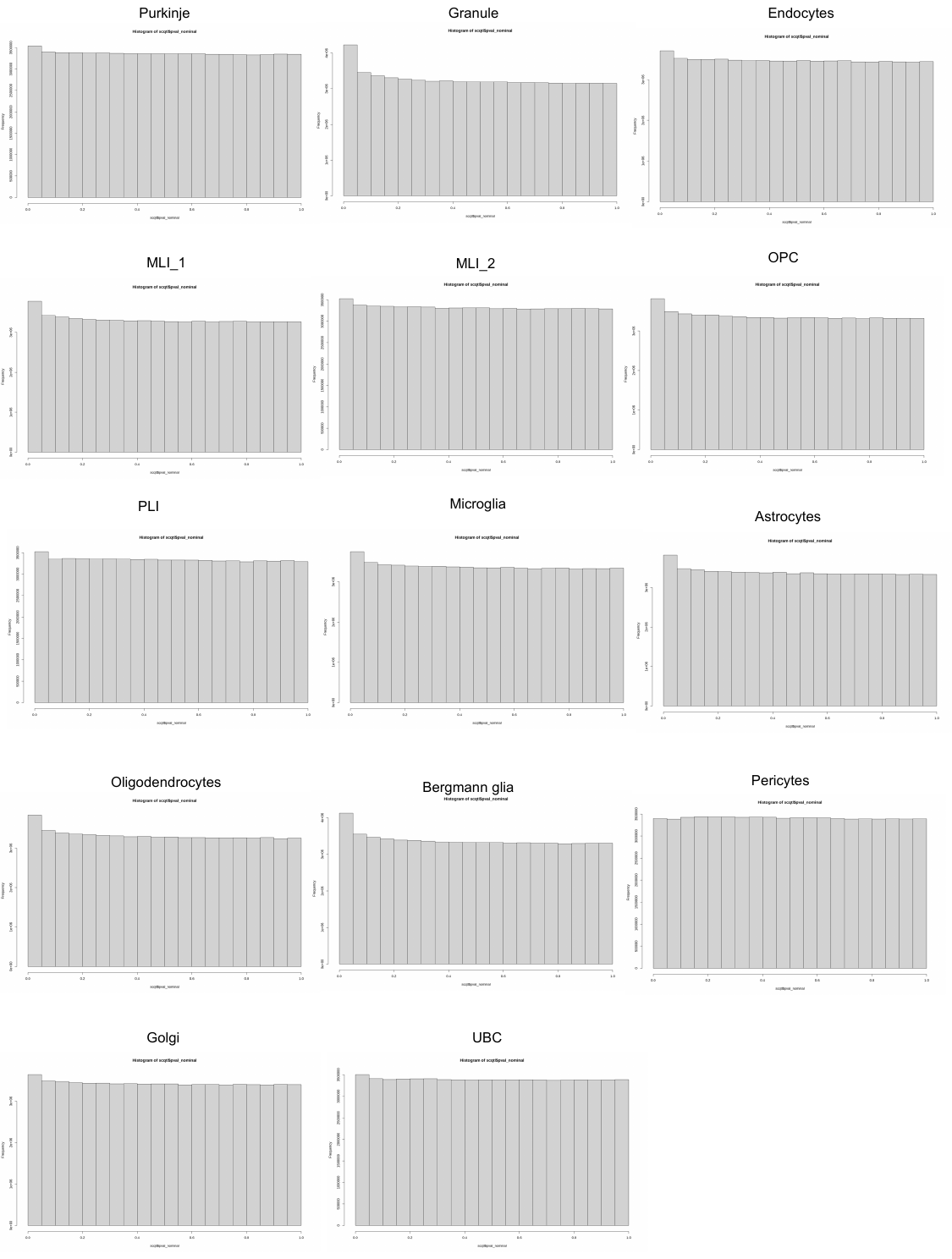

**Supplementary figure 2.** P-value histograms for cerebellar scQTLs per cell-types

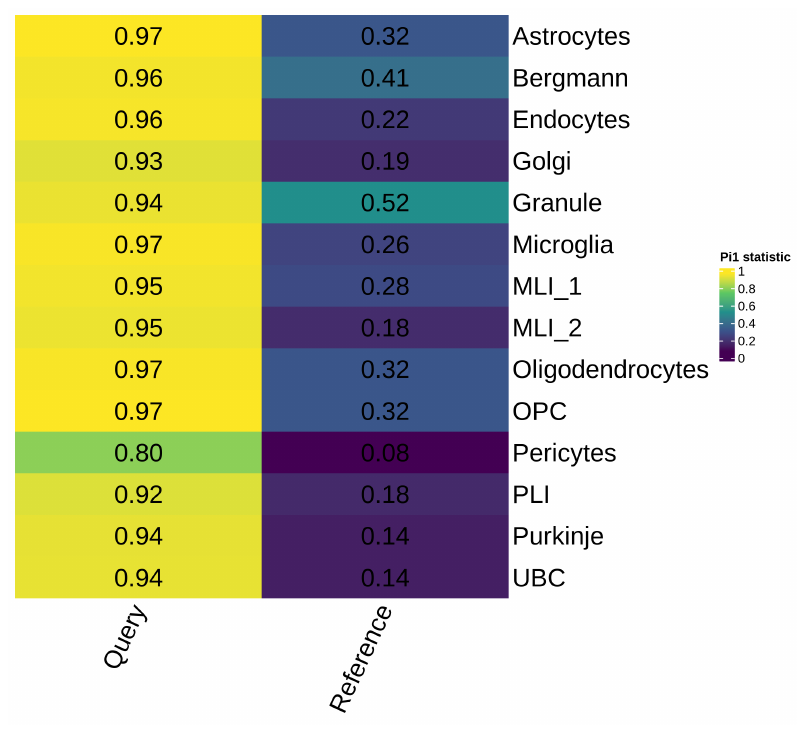

**Supplementary figure 3.** scQTL replication p-values with Cerebellum MetaBrain eQTLs.

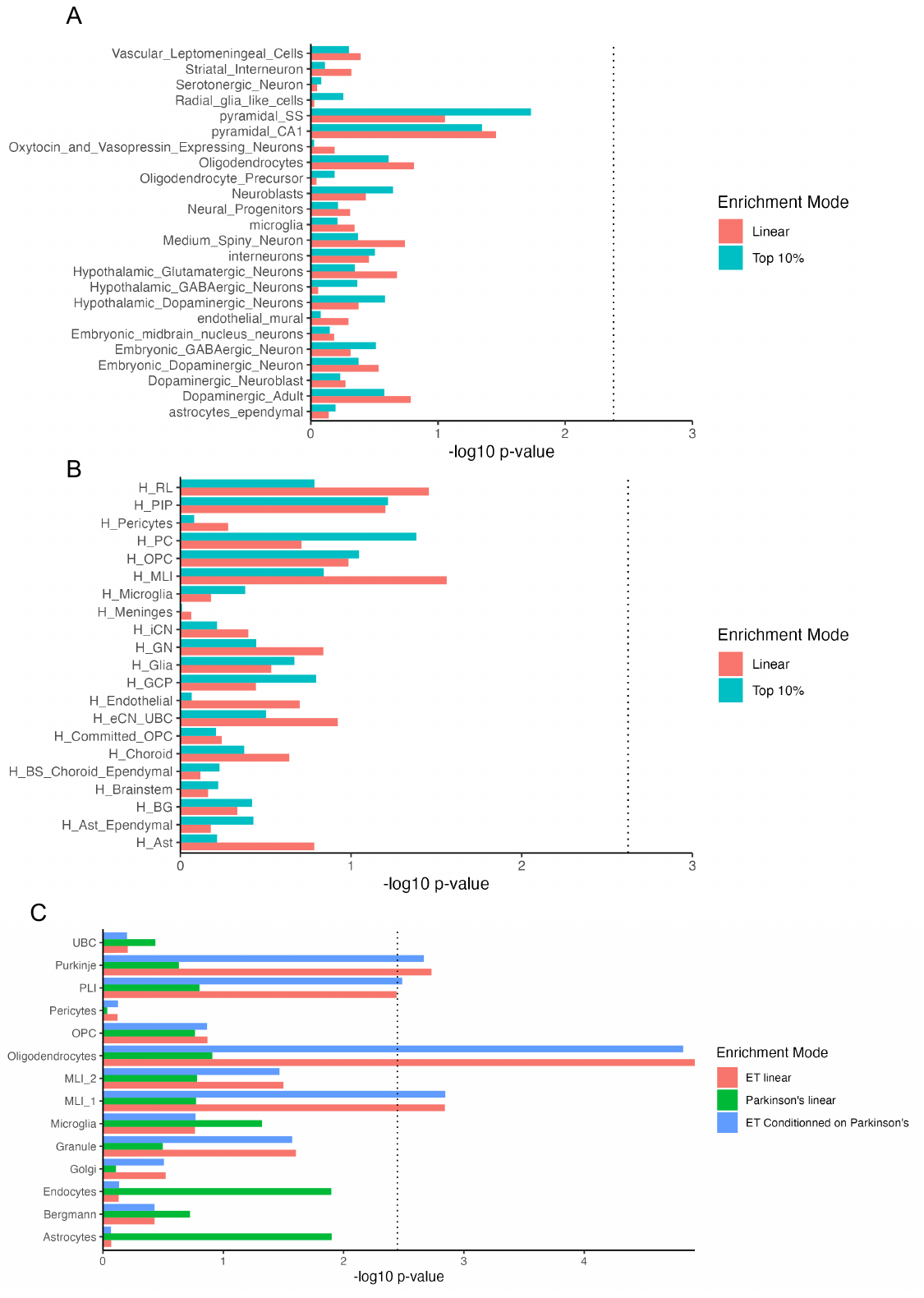

**Supplementary figure 4. MAGMA cell-typing.** A. ET GWAS enrichment in fetal cerebellum. B. ET GWAS enrichment in adult M1 cortex. C. PD genetic enrichment in cerebellar snRNA-seq.

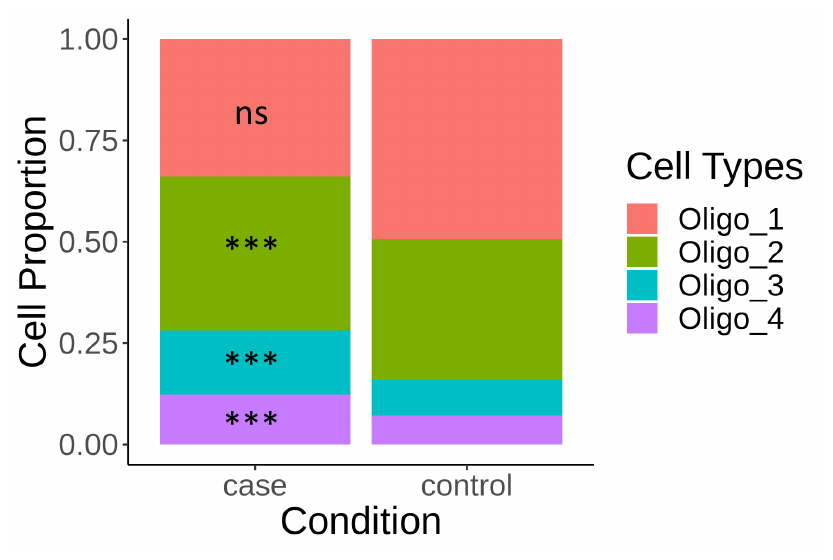

**Supplementary figure 5. Oligodendrocyte subclusters differences between ET and controls**. Oligo_1 adjusted p–value = 1.55e-1. Oligo_2 adjusted p-value = 2.41e-06. Oligo_3 adjusted p-value = 5.47e-8. Oligo_4 adjusted p-value = 2.57e-08. P-values were obtained from permutation testing without replacement (10,000 randomizations). **Legend:** ns = non-significant, * = 0.05, ** = 0.005, *** = 0.0005.

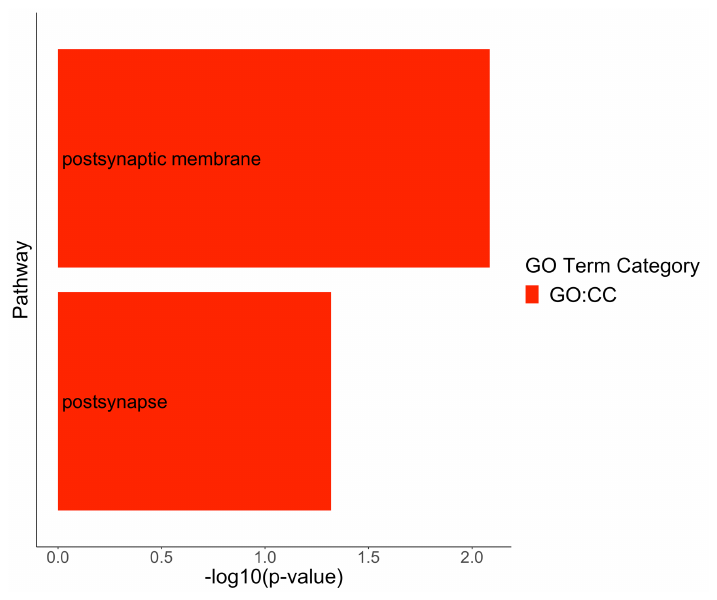

**Supplementary figure 6. Purkinje cells DEGs pathway enrichment analysis.**

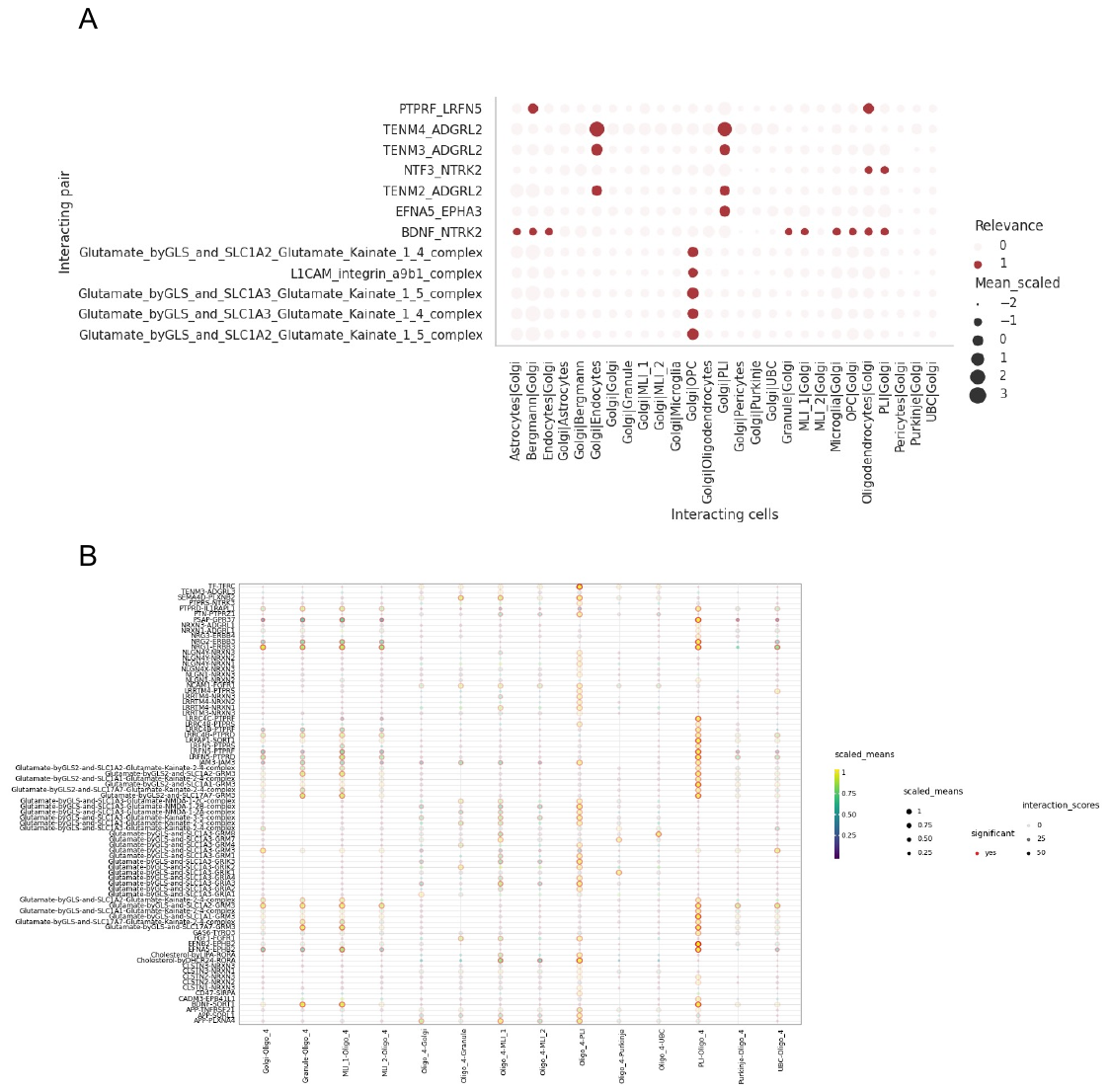

**Supplementary figure 7. CellphoneDB cell-cell communication analyses. A.** DEG-based analysis for cell-cell communications between Golgi cells and other cerebellar cells. Dot size indicates scaled expression of receptor-ligand complex across communicating cells. Relevance color indicates whether one or both signalling complexes are DEGs in either of the cell pairs. **B.** Oligo_4 interactions with other cerebellar cells using the statistical analysis method implemented in cellphoneDB. Size and color of dot indicates scaled expression of the receptor-ligand in cell pairs. Red contour indicates whether receptor-ligand interaction is significant (p-value < 0.05). Transparency indicates whether the interaction is specific to a certain cell pair. The higher the specificity, the higher the score.
